# Characterization of oral microbiota in marmosets: feasibility of using the marmoset as a human oral disease model

**DOI:** 10.1101/462911

**Authors:** Sachiko Takehara, Jorge L Zeredo, Yasuhiro Kumei, Kensuke Kagiyama, Kazumasa Fukasawa, Akiko Oshiro, Masayuki Ueno, Shunsuke Minakuchi, Yoko Kawaguchi

**Affiliations:** Department of Public Health, Tokyo Women’s Medical University, Tokyo, Japan; Department of Oral Health Promotion, Graduate School of Medical and Dental Sciences, Tokyo Medical and Dental University, Tokyo, Japan; Graduate Program in Health Sciences and Technology, University of Brasilia, Brazil; Department of Biochemistry, Graduate School of Medical and Dental Sciences, Tokyo Medical and Dental University, Tokyo, Japan; Clea Japan Inc., Tokyo, Japan; Gerodontology and Oral Rehabilitation, Graduate School of Medical and Dental Sciences, Tokyo Medical and Dental University, Tokyo, Japan

## Abstract

As the world’s population is aging, there is an increasing demand for research regarding aging and aging-related disorders, to achieve better understanding of aging. Ideally, such research would be performed with human subjects. However, due to ethical considerations, animals such as rodents and monkeys are used as alternatives. Among these alternative models, non-human primates are preferred because they share common traits with humans. The small South American common marmoset (*Callithrix jacchus*) may offer a number of advantages over other non-human primates in terms of its relatively small size, short life span, and identical dental anatomy, compared with humans. The purpose of this study was to clarify the viability of using the marmoset as a human oral disease model.

We collected saliva samples from eight common marmosets and eight human subjects. Prokaryotic DNA was extracted from the saliva samples, and 16S bacterial rRNA gene sequencing was performed on each of the samples. Our results indicated that the types of oral microbiomes detected among human and marmoset samples were nearly identical. In contrast, the oral microbiomes of our human and marmoset subjects were distinctly different from those of rat and dogs, which are popular animal models. The oral microbiomes of marmosets showed greater diversity than those of humans. With respect to individual variation, marmosets exhibited less variation in their oral flora, compared with humans. This difference in variation might be attributed to the fact that marmoset subjects were kept in a controlled environment with identical lifestyles.

The characteristics of its oral microbiota, combined with other technical advantages, suggest that the marmoset may provide the best animal model thus far for the study of oral health. This study characterized the oral microbes of the marmoset, thereby providing information to support future application of the marmoset as a model for age-related oral disease.

## Introduction

Aging is a complex phenomenon affecting all organs of the body. In the field of basic science, experimental subjects such as cells, worms, flies, and rodents have provided extensive groundwork for aging research [1]. However, taking this science from the bench to the bedside requires studies in more complex species with physiology and aging processes, which more closely resemble that of humans. Therefore, there is an increasing demand for research using non-human primates, which have higher phenotypic similarities and share a high genetic homology with humans [2]. The small South American common marmoset (*Callithrix jacchus*) offers a number of advantages over other non-human primates given their relatively small size and shorter life span [2]. Consequently, the marmoset has been used as a primate model of age-related diseases such as Parkinson’s disease, respiratory diseases, and infectious diseases.

As society rapidly ages, an increasing number of studies have been articulating the importance of maintaining oral health. One example is the association between the oral condition of elderly people and their quality of life in terms of susceptibility to aspiration pneumonia and falls [3]. In the adult marmoset, each oral quadrant contains two incisors, a short-tusked canine tooth, three premolars, and two molars, which totals 32 teeth [4]. In respect to teeth type and number, the oral anatomy of the marmoset is quite similar to that of humans.

More than 700 indigenous bacterial species inhabit the human oral cavity [5] and they form microbial communities within dental plaque and tongue coatings. Two major oral diseases, dental caries and periodontal disease, are infectious diseases closely related to the presence of specific oral bacteria. Bacteria in the human oral cavity have been extensively studied with great interest. However, bacteria from non-human primates have been studied less extensively. The purpose of this study was to examine the possibility of using marmosets as a human oral disease model. The present investigation was designed to elucidate the major bacteria in the marmoset’s oral cavity and examine the similarity with those in the human oral cavity using targeted sequencing of the microbial 16S rRNA gene.

## Materials and Methods

### Study subjects

Eight healthy common marmosets aged between 13 and 20 years were selected from the colony at Clea, Japan (Gifu, Japan). Human subjects were recruited among the patients who visited the Dental Hospital, Tokyo Medical and Dental University. Finally, eight healthy human subjects (male = 4, female = 4, age range: 37–61 years) were selected after excluding those with any prescribed medication, periodontitis, or systemic diseases.

### Saliva collection

Marmosets were fasted on the day of saliva collection in order to obtain undisturbed oral flora from each animal. Saliva samples were collected by probing sterilized cotton swabs (JCB Industry Limited, Tokyo) in the oral cavity and allowing the marmosets to chew on the cotton swabs habitually. After 2–3 minutes, the cotton swabs were removed and suspended in 1 ml of sterile distilled water in sterile 1.0-ml tubes. The tubes were then stored at –80 °C, until assayed for DNA analysis.

As for human subjects, they were asked to refrain from eating, drinking, brushing their teeth, and rinsing their mouth on the day of saliva collection in order to obtain undisturbed oral flora from each subject. Saliva was obtained by requesting them to spit saliva into sterile tubes while sitting in an upright position in a chair; saliva samples were frozen immediately and then stored at −80°C until analysis. In order to minimize the effects of circadian rhythm, samples from two species were collected between 9:00 a.m. and 11:00 a.m.

### Oral examination

All subjects underwent a standard oral examination. Oral examinations for marmosets and humans were conducted by registered dentists. Data such as types of teeth present, teeth condition (sound, decayed, or missing), and gingival inflammation were collected.

### DNA isolation and 16S rRNA gene sequencing analysis

DNA was extracted using a NucleoSpin Soil (Macherey-Nagel, Duren, Germany) according to the manufacturer’s instructions. The quantity and quality of extracted DNA were determined using a NanoDrop spectrophotometer (Thermo Fisher Scientific Inc., Japan), Quant-iT dsDNA HS Assay Kit (Thermo Fisher Scientific Inc., Japan), and agarose gel electrophoresis.

The fusion primers 341F (5’-TCGTCGGCAGCGTCAGATGTGTATAAGAGA CAGCCTACG GGNGGCWGCAG-3’) and 806R (5’-GTCTCGTGGGCTCGGAGATGTGTATAAGAGACAG GGACTACHVGGGTWTCTAAT-3’) with dual index were used for amplifying the V3–V4 regions of the bacterial 16S rRNA gene under the melting temperature of 50 °C with 28 cycles. The sequencing was performed on the Illumina MiSeq platform (Illumina, USA). Shannon diversity index, Chao diversity index, and unweighted UniFrac were calculated using QIIME (v. 1.8.0). The Man-Whitney test was used to compare the microbial composition between the two species.

For clustering of sequences into OTUs, CD-HIT-OTU (version 0.0.2) was utilized. Sequences were clustered into operational taxonomic units (OTUs) at 97% similarity using CD-HIT-OUT software. Taxonomic assignments for the 16S rRNA was performed by using RDP classifier program (version 2.2). A total of 51,135 sequences remained and were automatically used for clustering after quality control screening.

### Ethics statement

No animals were sacrificed during this study. All animals were cared for according to the protocol approved prior to the start of the study by the Animal Welfare Committee of Clea Japan Co. (Tokyo, Japan). All study animals were kept on a 12-hour light/dark cycle (7 a.m./7 p.m.) and were housed in isolated cages individually. The size of the cage was as follows: length 39 cm, width 55 cm, and height 70 cm. All were fed with commercially available monkey food for breeding and stud feed CMS-1M (FEED ONE CO., LTD., Japan) twice a day. Daily care was provided by the same staff, and veterinary staff took care of animals when they had any health problems. The veterinary staff also examined the conditions of animals before and after the study. As for the human subjects, the study was performed in accordance with the World Medical Association Declaration of Helsinki, and the study protocol and consent procedures were approved by the institutional ethics committee at Tokyo Medical and Dental University prior to commencement of the study (Approval No. D2015-606).

## Results

### Demographic characteristics

Human participants were given written and verbal explanation with respect to the purpose of this study. Eight subjects signed the consent forms and agreed to enroll in the study. As for animal subjects, eight healthy marmosets were randomly selected. Distribution of age among human subjects and marmosets is shown in Table 1.

**Table 1.** List of subjects and their characteristics.

| species | ID | age | gender | teeth<br>present | Number of OTUs |
| --- | --- | --- | --- | --- | --- |
| human |  |  |  |  |  |
|  | h-1 | 41 | Male | 28 | 105 |
|  | h-2 | 49 | Male | 27 | 132 |
|  | h-3 | 61 | Male | 25 | 91 |
|  | h-4 | 38 | Male | 28 | 124 |
|  | h-5 | 37 | Female | 24 | 75 |
|  | h-6 | 42 | Female | 28 | 104 |
|  | h-7 | 42 | Female | 24 | 98 |
|  | h-8 | 51 | Female | 20 | 84 |
| marmoset |  |  |  |  |  |
|  | m-1 | 10 | Male | 19 | 118 |
|  | m-2 | 12 | Male | 16 | 117 |
|  | m-3 | 15 | Male | 13 | 116 |
|  | m-4 | 18 | Male | 20 | 115 |
|  | m-5 | 12 | Female | 13 | 139 |
|  | m-6 | 12 | Female | 23 | 122 |
|  | m-7 | 15 | Female | 23 | 123 |
|  | m-8 | 12 | Female | 12 | 124 |

### Oral condition

The oral conditions compared are shown in Table 1. The number of present teeth among marmosets and humans varied from 12–23 and 20–28, respectively. Marmosets had a significantly smaller number of present teeth and a larger number of missing teeth than humans (p = 0.001). The main reason marmosets lost their teeth was habitual behavior such as chewing on hard wood. No subjects, in either human and marmoset groups, had decayed teeth.

### Saliva microbiome diversity

The distribution of the number of OTUs and genera are shown in Table 1. The number of OTUs was higher in humans than in marmosets (p = 0.016). In order to characterize the phylogenetic composition of bacterial communities in oral cavity, the α-diversity of microbiota were examined. The marmosets had a significantly higher Shannons index and Chao index than humans (Fig 1A, 1B, Mann-Whitney U test, p < 0.001, p = 0.0013), indicating a higher α-diversity of oral bacteria in marmosets.

**Fig 1.**
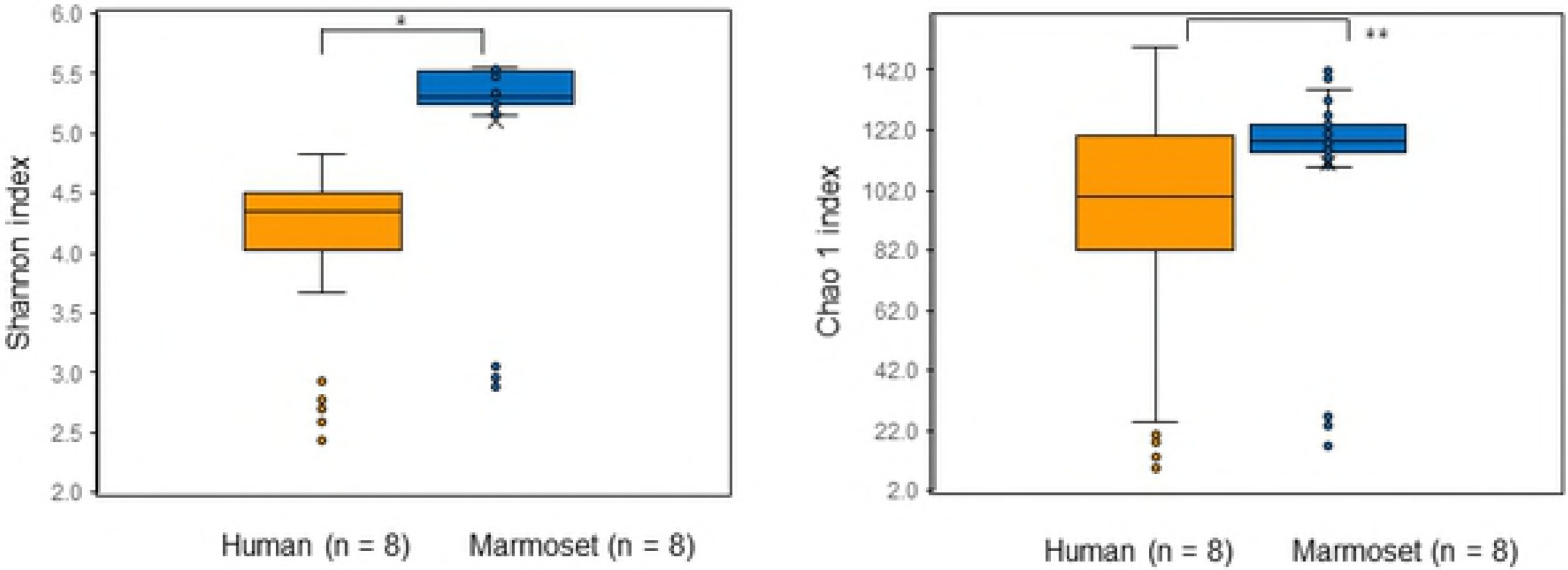
Diversity analysis for human and marmoset based on the bacteria abundance distribution for each individual at OTU level. (A) Shannon index diversity. *:p < 0.0001. (B) Chao 1 index diversity. **: p = 0.0013.

The overall bacterial community composition was compared using the UniFrac. UniFrac represents a phylogeny-based distance metric ranging from 0 (identical bacterial communities) to 1 (completely different). A principal coordinate analysis (PCoA) plot based on unweighted UniFrac values revealed distinct clustering of samples from marmosets and humans, indicating the difference in microbial compositions between the two species (Fig 2). As for their spatial distributions, samples from marmosets were densely clustered, whereas those from humans were loosely clustered. This tendency indicates that marmosets had smaller individual variations of their oral microbes.

**Fig 2.**
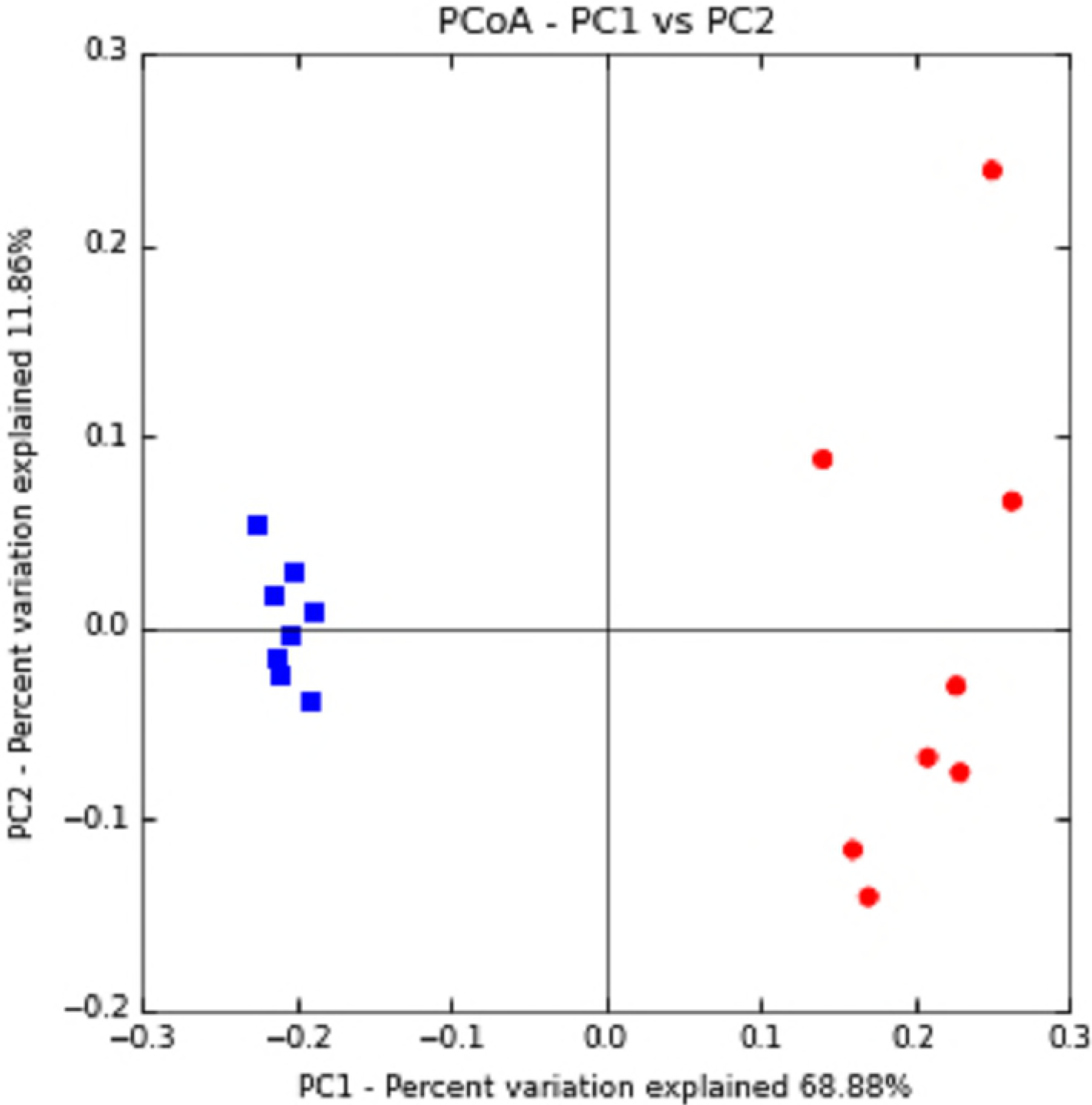
A principal-coordinate analysis plot of the oral microbiota based on the results of the unweighted UniFrac metric.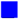: marmoset, 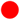: human

Furthermore, taxonomic distribution of the numerically abundant bacteria derived from the 16S rRNA gene sequences in all samples were examined (Fig 3). Most of the sequences obtained in this study were assigned to seven bacterial phyla—*Firmicutes, Bacteroides, Proteobacteria, Actinobacteria, Fusobacteria, TM7,* and *Spirohaetes.* The other phyla, Tenericues, Synergistetes, and SR1 were detected in the traceable few percent among the whole oral microbiome in our study samples. The average relative abundance of these seven phyla is shown in figure 4. *Firmicutes, Bacteroides,* and *Proteobacteria* were the major phylotype in human saliva, which accounted for 33%, 27%, and 21% of relative abundance on average, respectively. The forth abundant phylotype was *Actinobacteria*, which accounted for 9%. On the other hand, in the marmoset samples, *Proteobacteria, Bacteroides, Fusobacteria,* and *Firmicutes* were the major phylotype, which accounted for 26%, 24%, 21%, and 19%, respectively. Their relative abundance was statistically examined using the Mann-Whitney U test. Compared with humans, *Actinobacteria* and *Firmicutes* were statistically less abundant in marmosets (p = 0.002, p = 0.004), and *Fusobacteria* and *Spirochaetes* were statistically more abundant in marmosets (p < 0.001).

**Fig 3.**
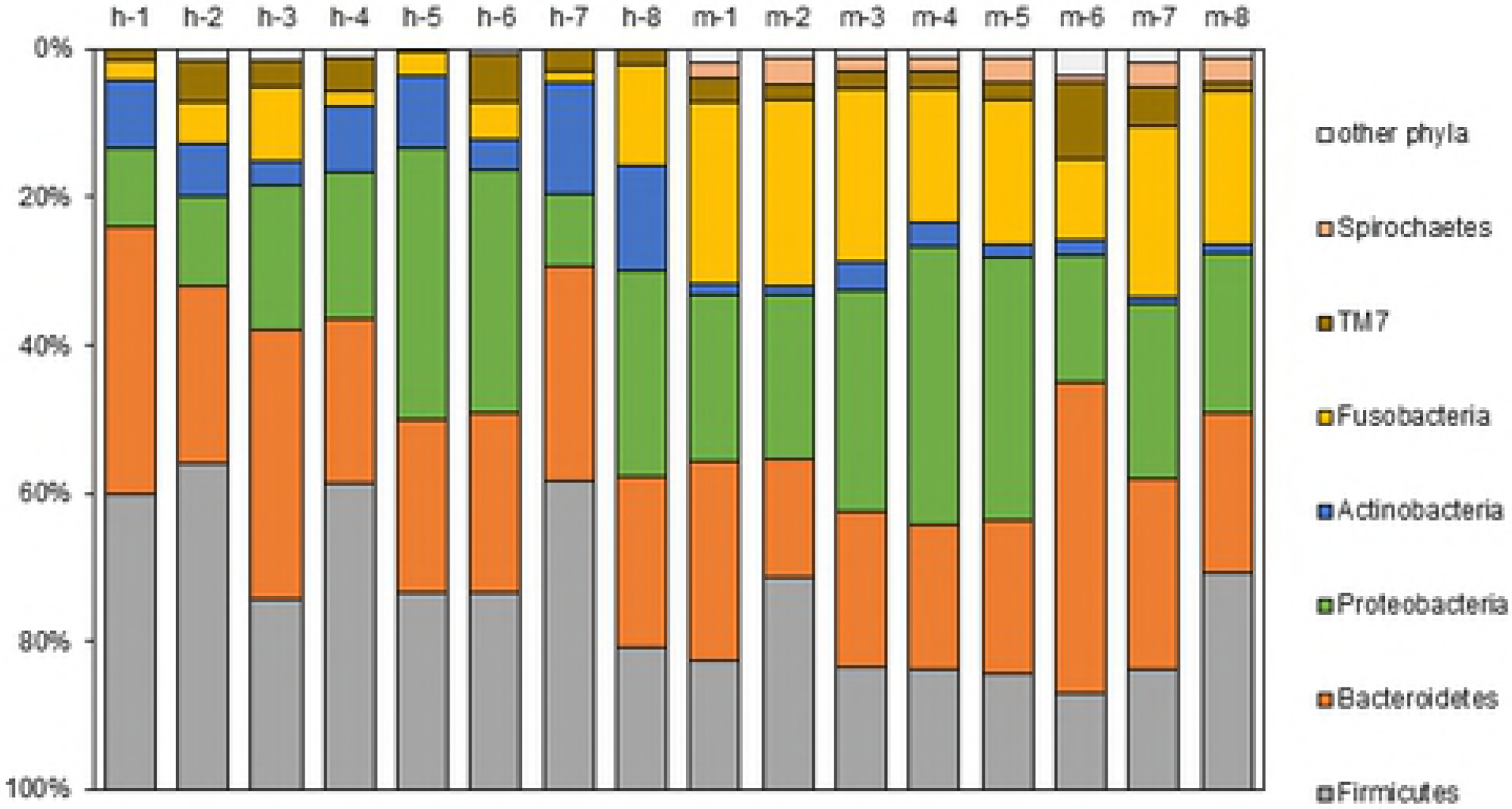
Microbial composition at the phylum level. Samples are represented along the horizontal axis, and relative abundance is denoted by the vertical axis. h-1 – h-8: human, m-1– m-8: marmoset.

**Fig 4.**
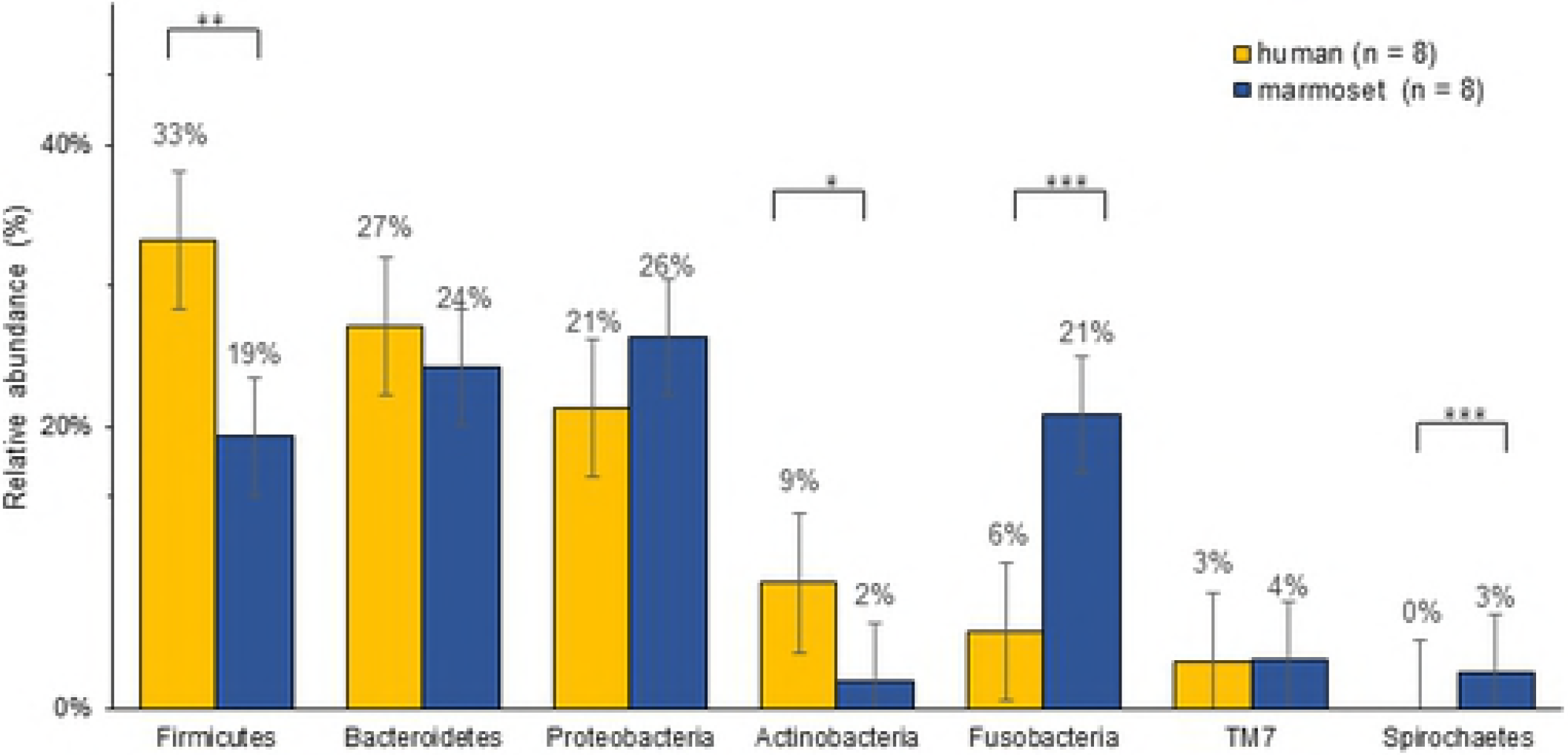
Relative abundance of Bacteria (Phylum level). *: p = 0.002, **: p = 0.004, ***: p < 0.0001

## Discussion

In this present study, we analyzed oral microbes from marmosets and humans for comparison. This is the first known report revealing the viability of marmosets as a non-human model for oral diseases and oral physiology. Our study has demonstrated that oral microbes in marmosets exhibit higher diversity with less individual variations than that of human oral microbes.

Anatomically, both humans and marmosets have common dental physiology in terms of number and types of teeth [4]. A marmoset’s life span is known to be between 16 and 21 years [6-8]—approximately one-seventh of a human life span. Such a life-span can be useful for intervention or observation studies investigating physiological and functional changes accompanied with aging. This versatility also can provide an opportunity for further studying the association between the elderly’s quality of life and poor oral health such as the increased incidence of aspiration pneumonia [3]. Aging is a multi-factorial phenomenon and no two individuals experience the same age-related decline. It is this relationship between the aging process and oral health that can be examined in the marmoset’s shorter life span, allowing us to observe the effect of poor oral health on the quality of life in a much more manageable time span.

This study revealed that oral microbe samples from marmosets showed higher diversity at the OTU level. Although, in general, oral microbiome varies dynamically over a life-time, this diversity is known to be maintained by healthy oral conditions through an optimal balance between the oral microbiota and the host immune system [9]. Any onset of periodontitis, dental caries, or even smoking may influence bacterial diversity [10]. In addition, this diversity decreases when mammal subjects experience chronic illness such as liver or gastrointestinal diseases [11, 12]. Both groups of our study subjects were healthy without any prescribed medication, diagnosed chronic diseases, or smoking habit. For marmosets, we selected those that were 12 years of age or older. With respect to their reported life-span [6], our marmoset subjects were categorized as being in old-age. Age, being one of the main risk factors for general prevalent diseases [13], brings susceptibility to infections by any compromised immune system. If this age factor reflects the host’s health condition [14], our senior-aged marmoset samples should have lost oral microbiome diversity. Quite contrary to this, we observed that they maintained a high microbial diversity.

There were a number of distinctive differences observed in the salivary microbiome between humans and marmosets. Previous studies reported clear differences among populations living under different geographical conditions [15, 16]. The differences were reportedly attributed mainly to differences in lifestyles, dietary habits, as well as sanitary and socioeconomic factors [17, 18]. Our human oral microbiome diversity data was consistent with the Human Oral Microbiome Database established by Delwhirst et al. [19]. In respect to individual variation, oral microbes of marmosets had less variety in their oral flora compared to human data, which exhibited higher variation. This difference in variation can be attributed to the fact that marmoset subjects were kept in a controlled environment with identical lifestyles (i.e. marmosets were kept in similar cages and fed with identical food). We had expected that the phylum *Spirochaetes*–reported to include species of the genus *Treponema*, found in root canal infections, gingivitis, and periodontitis in humans [19]–would not be detected in human samples. Conversely, *Spirochaetes* was detected in our marmosets’ samples because we did not select marmosets according to their oral-health status; our selection process was based strictly on their age group.

Lab rats, such as Wistar rats, and canines have been widely used in scientific experiments as models for human oral disease. The oral microbiomes of lab rats [20] and canines [21] have been reported. Oral microbiomes of canines are similar to those of humans, but still a significant gap of phylum difference exists. As for rats, distribution of oral microbiota is reported to be even more different than that of humans. In contrast, marmosets’ oral microbiome is much closer to that of humans, as compared to reported microbiota of the rat and canine.

A major limitation of this study was that our sample consisted exclusively of old-age marmosets. Although we selected marmosets of a wide age range (10 to 18 years), our sample turned out to be surprisingly homogeneous in terms of oral microbiome and oral-health status. Therefore, we could not make any associations between these factors (age, microbiome, and health/disease) within our sample. Nevertheless, it is reasonable to assume that such associations not only exist, but also that they can be teased out in future studies by including marmosets of younger age-groups and by collecting more detailed data on the marmosets’ oral health status, such as the severity of gingival inflammation, depth of periodontal pockets, and extent of dental decay. Such studies shall reinforce the feasibility of the marmoset as an ideal, alternate oral health model for future research.

In the context of the marmoset’s dental physiology, shorter life span, and smaller body mass, our findings strongly suggest that the marmoset could be an ideal model for studying general oral health and its relationship with aging. In conclusion, this study is the first to elucidate the oral microbes of the marmoset’s, thereby providing important information for the application of the marmoset as a model in studies of age-related oral diseases.

## Acknowledgements

We are grateful to Clea Japan for supporting our study by offering marmosets and space to conduct our study.

